# Antigenicity and Potential Use of A Novel Brucella Multiepitope Recombinant Protein in the Diagnosis of Brucellosis

**DOI:** 10.1101/786343

**Authors:** Dehui Yin, Qiongqiong Bai, Li Li, Lichun Xu, Kun Xu, Juan Li

**Author notes:** Equal contributors (Dehui Yin, Qiongqiong Bai). Correspondence: Lichun Xu, School of Public Health, Xuzhou Medical University, No. 129 Tongshan Road, Xuzhou, 221004, China. & Kun Xu, School of Public Health, Jilin University, No. 1163 Xinmin Street, Changchun, 130021, China.

## Abstract

Currently, brucellosis is a reemerged zoonotic infectious disease with an increased incidence in recent years. A simple, rapid and sensitive method for diagnosing brucellosis has important significance for early diagnosis and early treatment, which can help to reduce medical burden and economic loss. Previously, a multiple epitope recombinant protein was constructed based on linear B-cell epitope prediction tools. In this study, the recombinant protein was used as an antigen to study the immune response produced by immunized mice, and goat serum was used to verify its diagnostic accuracy.The production of antibodies was successfully induced in the vaccinated mice. Through analyzing the serum antibody subtypes, the primary antibody was identified as IgG. Flow cytometric analysis revealed that the percentage of CD4+, CD8+ and the CD4+/CD8+ ratios were increased by T cell subsets in mouse splenocytes, indicating that the recombinant protein induced a strong immune response in mice and that it had strong immunoreactivity. Using indirect ELISA, the recombinant protein correctly diagnosed positive and negative brucellosis samples. Compared with the whole bacterial antigen, the recombinant protein had a weaker sensitivity but a stronger specificity.In this study, animal experiments showed that the recombinant protein had good antigenicity, and indirect ELISA indicates that it can be used as an antigen to diagnose brucellosis. Therefore, the recombinant protein is a potential candidate antigen for brucellosis vaccine development and serological diagnosis.

## Introduction

Currently, a high incidence of brucellosis, a hazardous zoonosis, is reemerging worldwide, especially in developing countries. There are more than 500,000 new cases every year in the world[1], and the number of cases in China is also increasing annually, with approximately 37,000 new cases reported in 2018 (http://www.nhc.gov.cn/jkj/s3578/201904/050427ff32704a5db64f4ae1f6d57c6c.shtml). Brucellosis causes not only human health problems but also extensive losses in agriculture and livestock husbandry. Therefore, it is of great importance to strengthen the prevention and diagnosis of brucellosis.

In recent years, various diagnostic methods based on detecting specific antibodies in patient sera have been used for the diagnosis of brucellosis. The Rose Bengal plate test (RBPT), plate agglutination test (PAT), standard tube agglutination test (SAT), complement fixation test (CFT), Coombs anti-Brucella test and enzyme-linked immunosorbent assay (ELISA) are the main methods for the serological diagnosis of brucellosis. However, these methods usually use the whole cell or smooth lipopolysaccharides (S-LPS) as the antigen to detect *Brucella* antibodies in the patient’s serum, which can lead to false positives and cross-reactivity. The existing brucellosis vaccines have some potential safety hazards. Additionally, most brucellosis diagnostic methods are time-consuming, of low-sensitivity, and cross-reactive[2,3]. The key to improving brucellosis prevention and diagnosis is to find suitable antigens. Therefore, the development of new antigens is crucial for improving the safety of brucellosis vaccines and the specificity and sensitivity of diagnostic methods. A large number of studies in immunology have shown that *Brucella* outer membrane proteins (OMPs) are very immunogenic, and they are potentially new subunit vaccine and diagnostic antigens for brucellosis[4,5]. With the rapid development of computer science and the large-scale development of genome and proteome sequencing, a new discipline has emerged called bioinformatics. Bioinformatics is an interdisciplinary subject that integrates computer science, life sciences, and mathematics. It is widely used in genomics and proteomics. In terms of proteomics, bioinformatics can predict the structure and function of proteins and has been used for predicting protein structure and B cell and T cell epitopes[6,7].

In previous work, a new *Brucella* multiepitope fusion protein was successfully designed using the predominant B-cell epitopes of the main OMPs of *Brucella* and a new *Brucella* recombinant multiepitope protein (rMEP) containing B-cell epitopes of *Brucella* OMP16, OMP2b, OMP31 and BP26 was successfully designed using bioinformatics-related technology[8]. Based on previous work, the immunogenicity and antigenicity, the ability to recognize *Brucella* serum antibodies and the potential applications of the recombinant protein were assessed and evaluated in this study.

## Material and methods

### Animals and ethics statement

A total of 20 female SPF BALB/c mice, 6-8 weeks, weighing 18±2.0 g, were purchased from Beijing Hua Fukang Biological Polytron Technologies Inc. After the study, all mice were euthanized by CO_2_. All animal experiments were fully compliant with an ethical approval granted by the Animal Care and Ethics Committee of Jilin University (permit number: LSXK2015101) and the Institutional Research Ethics Committee of Medicine, the School of Public Health, Jilin University (permit number: JLU2014-0303).

### Serum samples

A total of 93 goat serum samples included 57 brucellosis-positive sera (*Brucella melitensis*, *B. melitensis*) that were determined serologically positive by plate agglutination tests (PATs) and standard tube agglutination tests (SATs), which are standard methods for diagnosing brucellosis, and 36 brucellosis-negative sera. All samples were provided by the China Animal Health And Epidemiology Center (Qingdao, China).

### Identification of immunogenicity of rMEP

#### Mouse vaccination

Before the start of the experiment, the BALB/c mice were kept in cages with a 12-h light-dark cycle and with water and food *ad libitum* for 1 week. BALB/c mice were randomly divided into two groups consisting of 10 mice per group. Group 1 was immunized by intraperitoneal (i.p.) injection of 25 μg of rMEP, and group 2 was intraperitoneally injected with phosphate-buffered saline (PBS). Antigens and PBS were mixed with equal volumes of complete Freund’s adjuvant (Sigma-Aldrich, St. Louis, MO, USA) on day 0 and with equal incomplete Freund’s adjuvant (Sigma-Aldrich) on days 15, 30, and 45. Sera were obtained before each immunization and stored at −70°C until assayed.

#### Humoral immune response

As described previously[9], enzyme-linked immunosorbent assay (ELISA) was performed to measure total IgG, IgM, IgG1, IgG2a, IgG2b, and IgG3 levels of antigen-specific serum according to manufacturer instructions (Southern Biotech, Birmingham, AL, USA), and 96-well plates (Corning, NY, USA) were coated with 100 μL/well purified protein [final concentration 10 μg/mL in PBS (pH 7.4)] and incubated overnight at 4°C. Wells were washed three times with PBS containing 0.05 % Tween-20, and plates were blocked with PBS containing 1 % bovine serum albumin for 2 h at room temperature (RT). The plates were later incubated with 100 μL of mouse sera at a 1:100 dilution for 1 h at RT and washed three times as described. Horseradish peroxidase (HRP)-labeled goat anti-mouse IgG, IgM, IgG1, IgG2a, IgG2b, and IgG3 (100 μL/well) were added and incubated for 1 h at RT. After a final wash step, 0.1 mL of 2,2-azino-bis(3-ethylbenzothiazoline-6-sulfonic acid) (ABTS) [15 mg ABTS powder dissolved in 1 mL of distilled water and diluted with 9 mL of substrate buffer consisting of 2.1 g of citric acid dissolved in 100 mL of H_2_O and 10 μL of 30 % hydrogen peroxide (pH 4.0)] substrate solution was added to each well. The optical density (OD) at 405 nm (OD405) of each well was recorded at 10 min and 20 min after substrate addition.

#### Flow cytometry analysis of lymphocyte subtypes

Splenocytes (100 μL; 1 × 10^6^ cells/mL) were stained with allophycocyanin-cyanine 7 (APC-Cy7)-labeled anti-mouse CD3, fluorescein isothiocyanate (FITC)-labeled anti-mouse CD4, and phycoerythrin (PE)-labeled anti-mouse CD8 (Invitrogen, USA). FITC-, PE-, and APC-Cy7-conjugated primary antibodies were used as isotype controls. Based on their scattering profiles, lymphocytes were gated, and the percentages of CD3-, CD4-, and CD8-positive cells were quantified. Data analysis was performed using NovoExpress software (Acea Biosciences, San Diego, CA, USA).

### Identification of the diagnostic accuracy of rMEP

To verify the sensitivity and specificity of rMEP in the diagnosis of goat brucellosis, an indirect ELISA assay was performed as described previously[8]. Meanwhile, the antigen in the SAT (the whole bacteria antigen was supplied by China Animal Health And Epidemiology Center) was used to evaluate the effectiveness of the rMEP protein as a diagnostic antigen for brucellosis by indirect ELISA. The whole bacterial antigen (dilution with 1:160) and the rMEP protein (100 μg/mL) were used as antigens in coating buffer (0.01 M PBS, pH 7.4) and incubated in 96-well microtiter plates (Corning) overnight at 4 °C. After washing 4 times with PBST (PBS containing 0.05 % Tween-20), the plates were blocked with 0.1 % ovalbumin (OVA, TCI, Japan) for 1.5 h at 37 °C, washed with PBST 4 times, and then incubated with a 1:400 dilution of goat serum at 37 °C for 60 min. After washing 4 times with PBST, the plates were incubated with HRP-conjugated rabbit anti-goat (Abcam, UK) immunoglobulin IgG antibodies at 1:10,000 dilutions for 1 h at 37 °C. The plates were washed and 100 μL TMB (Trimethylbenzene) substrate solution was added to each well. Reactions were left for 15 min at RT in the dark. Finally, 50 μL of stop solution (2 M H_2_SO_4_) was added to each well. Optical density values were measured at 450 nm using an ELISA plate reader (BioTek). All samples were tested in replicates.

### Statistical Analysis

Dot plot and receiver-operating characteristic (ROC) analyses were performed using GraphPad Prism version 6.05 for Windows. The OD405, OD450 and the percentages of lymphocyte subtypes were determined by Student’s t-test (unpaired t-test). Differences were considered statistically significant when *p* < 0.05.

## Results

### Immunogenicity of rMEP

After mice were immunized with rMEP, the serum titer of the mice rapidly increased compared with the control group, indicating that the recombinant protein used in the present study had good immunogenicity as shown in Fig. 1. Further analysis of antibody subtypes in serum showed that the main antibody subtypes were IgG, IgG1, IgG2a, IgG2b, and IgG3, which all had higher OD values, as shown in Fig. 2. The analysis of T cell subtypes in mouse spleen cells showed the experimental group had significantly higher CD4 differentiation than the control, and the CD4+/CD8+ ratio was increased, indicating that the recombinant protein significantly increased the Th1 response in mice. The recombinant protein has good immunogenicity, as shown in Fig. 3 and Table 1.

**Fig. 1.**
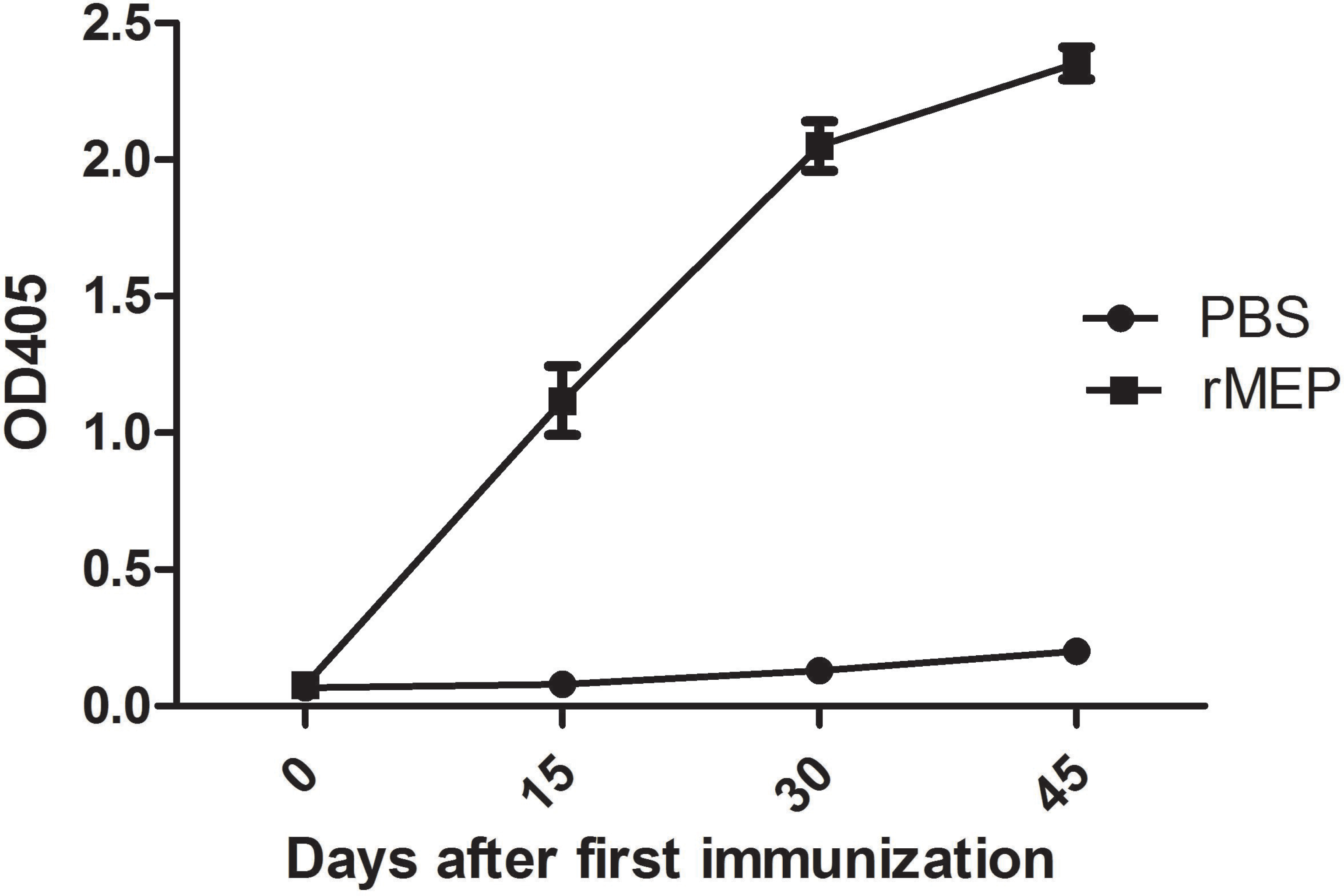
Changes in serum titer levels after immunization in mice. Test: rMEP; Control: PBS

**Fig. 2.**
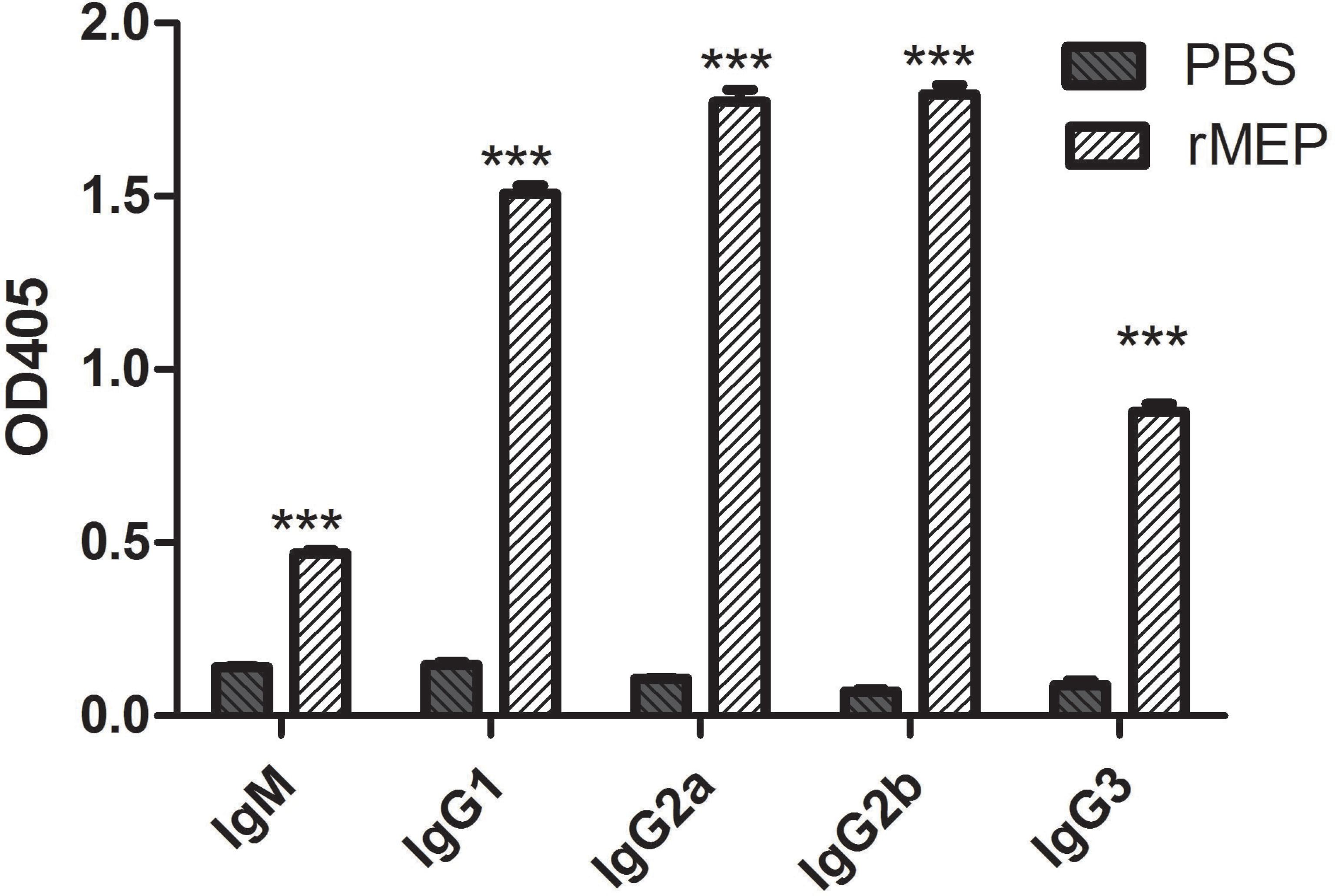
Antibody subtypes in mouse serum. Test: rMEP; Control: PBS; ***p<0.001

**Fig. 3.**
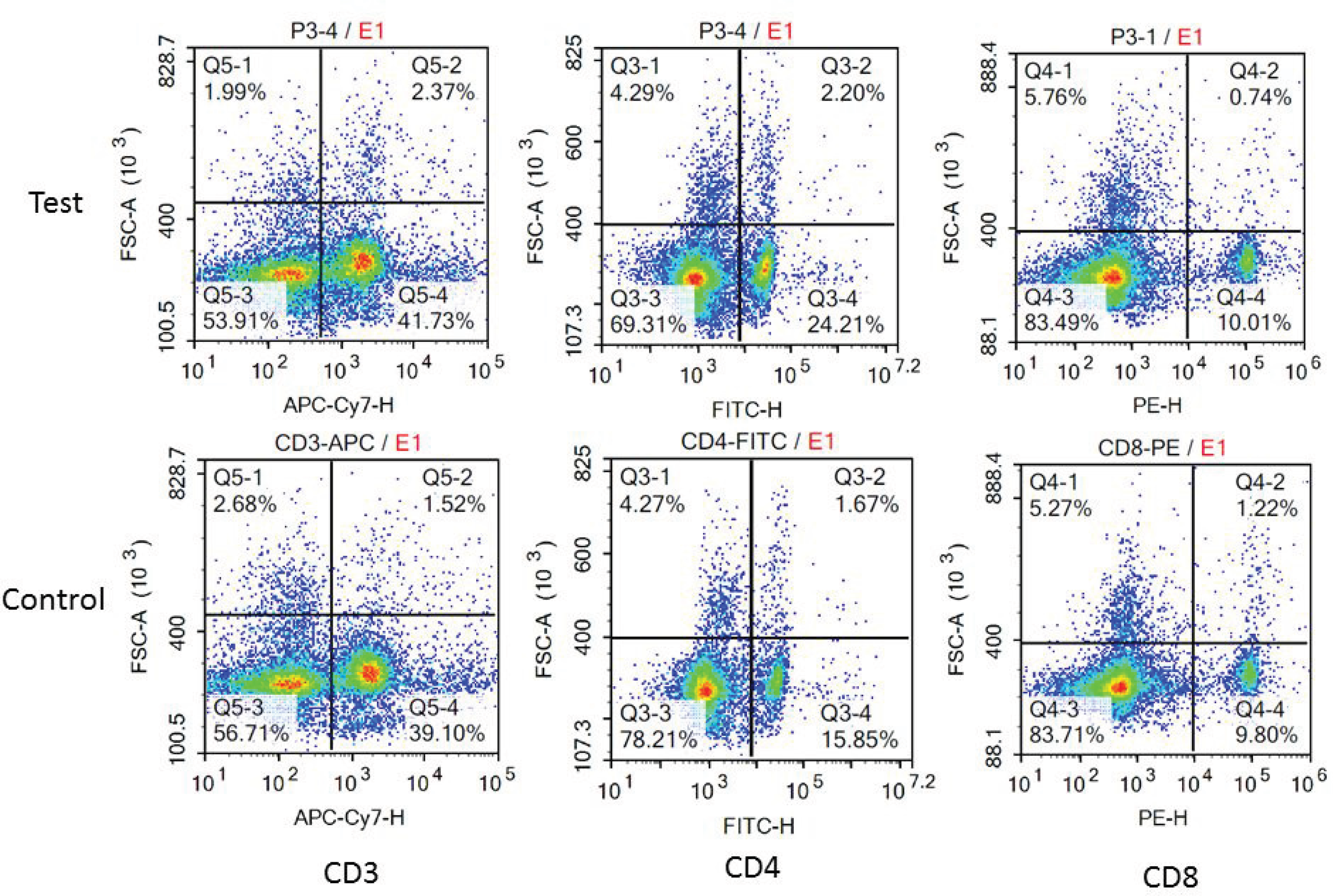
Flow cytometric analysis results. Test: rMEP; Control: PBS

**Table 1.** Percentage of CD3+, CD4+, CD8+ and CD4+/CD8+ by flow cytometric analysis (%)

| Group | CD3+ | CD4+ | CD8+ | CD4+/CD8+ |
| --- | --- | --- | --- | --- |
| Test | 44.69±0.81* | 27.57±0.99* | 10.94±0.14 | 2.54±0.059* |
| Control | 40.54±0.27 | 20.17±2.75 | 10.19±0.39 | 2.08±0.302 |

\*Compared with the control group, $p<0.05$

### Diagnosis accuracy of rMEP

To evaluate the accuracy of the rMEP for diagnosis of brucellosis, 93 serum samples were tested by indirect ELISA. A dot plot diagram (Fig. 4a) and receiver operating characteristic (ROC) curve analysis (Fig. 4b) were performed to evaluate the sensitivity and specificity. Based on the ROC curve analysis, the area under the curve (AUC) for rMEP was 0.9798 (95 % confidence interval (CI), 0.9569 to 1.003). In addition, a diagnostic sensitivity of 96.49 % (95 % CI, 8789 to 99.57) and a specificity of 94.44 % (95 % CI, 81.34 to 99.32) were obtained from an optimum cutoff value (0.6195). With this cutoff value, 55 of 57 brucellosis cases were diagnosed correctly as positive, and 34 of 36 negative cases were diagnosed correctly as negative. Furthermore, SATs using the whole bacteria antigen were used to diagnose brucellosis in parallel with indirect ELISA with the rMEP, and a dot plot diagram (Fig. 4c) and ROC curve (Fig. 4d) were constructed. The AUC was 0.9484 (95 % CI, 0.9040 to 0.9927), and a diagnostic sensitivity of 100.0 % (95 % CI, 93.73 to 100.0) and a specificity of 77.78 % (95 % CI, 60.85 to 89.88) were obtained from an optimum cutoff value (0.8875). With the cutoff value, all brucellosis cases were diagnosed correctly; however, only 28 of 36 negative cases were diagnosed correctly. A cross-table with absolute numbers of positive and negative samples with these cutoff values is listed in Table 2. The results of the ELISA indicate that it is possible to clearly and correctly differentiate between positive and negative brucellosis samples with this assay. Compared with the whole bacterial antigen, rMEP has a weaker sensitivity but a stronger specificity.

**Table 2.** Positive and negative predictive values of different cutoff values

| Cutoff value | positive |  | negative |  | PPV(%) | NPV(%) |
| --- | --- | --- | --- | --- | --- | --- |
|  | TP | FN | TN | FP |  |  |
| 0.611<sup>a</sup> | 55 | 2 | 34 | 2 | 96.49 | 94.44 |
| 0.887<sup>b</sup> | 57 | 0 | 26 | 8 | 87.69 | 100 |

TP, true positives; TN, true negatives; FP, false positives; FN, false negatives; PPV, positive predictive value $(TP/(TP + FP)) \times 100$ ; NPV, negative predictive value $(TN/(TN + FN)) \times 100$
a, cutoff value is calculated by rMEP; b, cutoff value is calculated by antigen of SAT

**Fig. 4.**
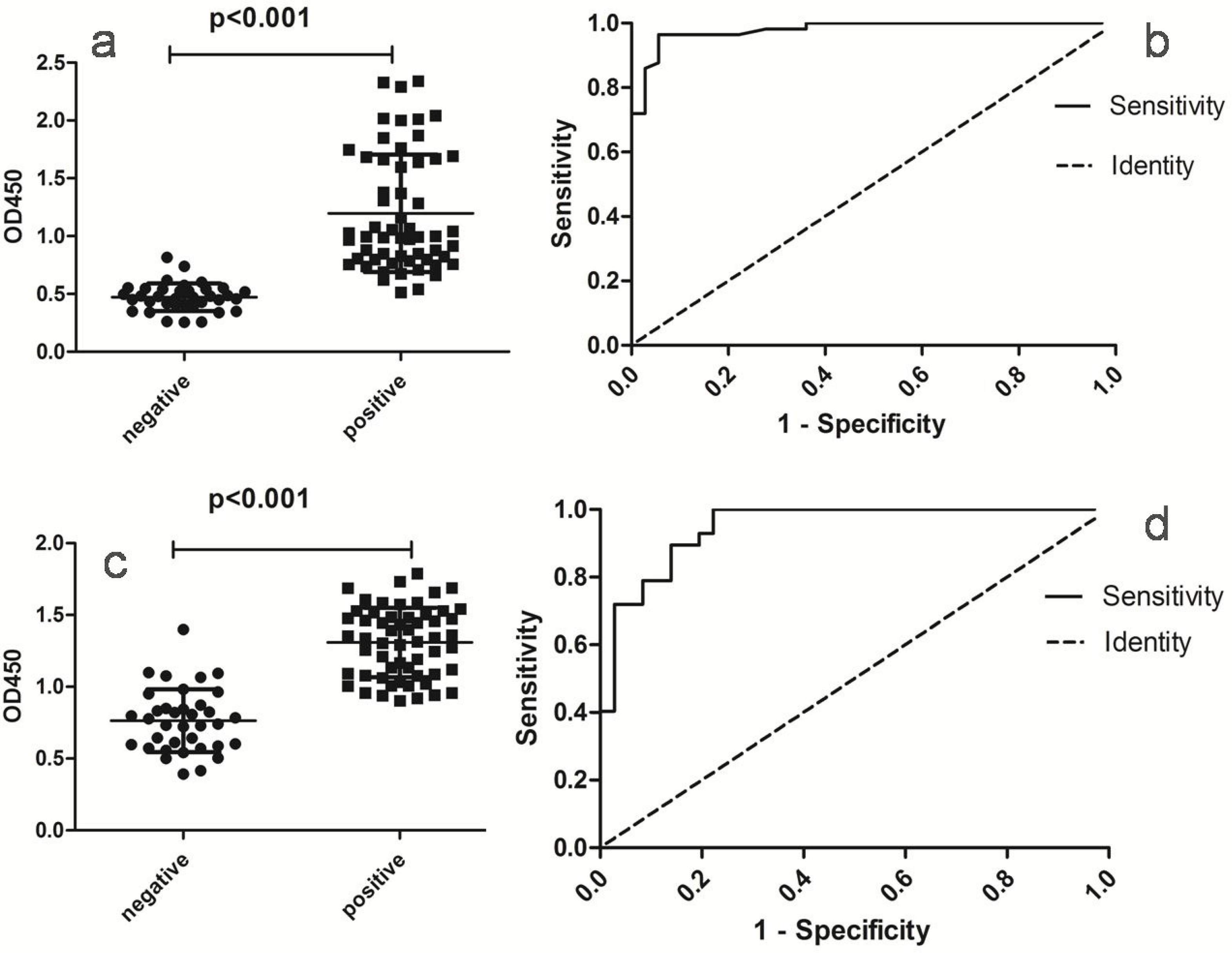
Indirect ELISA analysis of serum samples. The analysis was performed using positive brucellosis (57 sera) and negative control (36 sera) sera from goats. (a) Dot plot of the rMEP, (b) ROC analysis of rMEP, (c) Dot plot of the SAT antigen, (d) ROC analysis of SAT antigen

## Discussion

At present, many vaccine studies have shown that many proteins or other components of *Brucella* have strong antigenicity. Animal experiments show that these components have a certain degree of immunoprotection against *Brucella* and are good candidates for vaccine development. Therefore, these components also provide direction for researchers to develop new brucellosis diagnostic antigens, and the development of immunoinformatics technology provides tools for the development of new diagnostic antigens. Serological methods are the most commonly used methods for diagnosing brucellosis, but serological diagnostic methods have some disadvantages, such as low sensitivity and specificity. Thus, improving the diagnostic accuracy of antigens is key to improving the diagnostic sensitivity and specificity of serological methods. Specificity using *Brucella* OMPs in the diagnosis of brucellosis has certain advantages, but the sensitivity of a single OMP as an antigen for the diagnosis of brucellosis is restricted[10]. Combining multiple OMPs can significantly improve the diagnostic accuracy and increase the sensitivity[11]. To ensure its diagnostic accuracy, the rMEP contained B-cell epitopes of *Brucella* OMP16, OMP2b, OMP31 and BP26. Recombinant proteins are different from native proteins and may lose some function after expression. Therefore, after purification of the rMEP protein, relevant experiments were performed to verify its validity. The mouse model is the most commonly used to verify whether fusion proteins are immunogenic or antigenic[12,13,14]. The fusion protein constructed in this study was composed of the predicted linear epitopes of B-cells and was used to immunize mice. The antibodies were produced in the serum of the immunized mice and verified by indirect ELISA.

Antibody levels against rMEP specifically were measured in sera obtained 45 days after the first immunization. Antibody titer results demonstrated that antibodies in mouse serum were produced by rMEP. Further analysis of the antibody subtypes found that IgG was the main antibody produced in response to *Brucella* infection, accounting for more than 90 % of all antibodies[15]. The analysis of mouse T cell subtypes showed that the ratio of CD4+/CD8+ was significantly higher and the fusion protein was very immunogenic. After immunization, both strong humoral and cellular immune responses were induced in mice. Vaccine candidates should stimulate cell-mediated immunity, such as the induction of Th1 and Th2 immune responses, which are extremely important in controlling intracellular organisms such as *Brucella*. However, one of the major flaws in this study was the deficiency of in-depth research on cell-mediated immunity, such as cytokine secretion measurements (IFN-γ and IL-6), so the specific potential applications of rMEP should be considered cautiously.

In China, the government requires serological methods such as PAT, SAT, and iELISA for primary screening, and a general positive result will initiate eradication of the positive livestock or the whole farm, so SAT or iELISA is still used in China.

Although the current golden test of diagnosis is the pathogen culture, the positive rate of culture is too low, and the government does not mandate the identification of pathogens as long as the serological positive results are available, which is related to China’s strict management strategy. According to the epidemiological study of brucellosis, *Brucella* spp., including *B. melitensis*, *B. abortus*, *B. canis*, and *B. suis*, can infect humans and domestic animals[16], and human *B. melitensis* is the main *Brucella* species worldwide, which accounts for 80 % to 90 % of the total number of brucellosis cases in China[17,18]. Therefore, brucellosis sera infected by *B. melitensis* were used in this study to further verify the diagnostic accuracy of the recombinant protein. The results of indirect ELISA showed that the recombinant protein had good ability to recognize the positive- and negative-brucellosis sera. To investigate the sensitivity and specificity of the rMEP, ROC curve analysis was performed, and the results indicated that the rMEP had better specificity than the whole bacterial antigen. The AUC indicated that both the recombinant protein and the whole bacterial antigen could distinguish positive samples from negative samples with a high level of accuracy (0.9 < AUC < 1)[19]. The following work should be to expand the range of strains to further verify their specificity.

## Conclusion

The recombinant protein was used to immunize mice and successfully induced the production of antibodies in mice, resulting in a strong immune response, demonstrating that the recombinant protein had good immunogenicity. The indirect ELISA results proved that the recombinant protein had good sensitivity and specificity for diagnosing brucellosis in goat serum. It is a potential candidate antigen for brucellosis vaccine development and serological diagnosis.

## Declaration of conflicting interests

The authors declared no potential conflicts of interest with respect to the research, authorship, and/or publication of this article.

## Authors’ contributions

D-HY, Q-QB, KX and JL conceived and designed the study. D-HY, Q-QB and LL performed the animal and ELISA assays. D-HY, Q-QB and LL analyzed the data. D-HY drafted the manuscript. L-CX, JL and KX reviewed and made improvements in the manuscript. All authors read and approved the final version of the paper.

## Funding

This work was supported by the Natural Science Foundation of the Jiangsu Higher Education Institutions of China (Grant number 18KJB230005), Young Scientists Fund of the National Natural Science Foundation of China (Grant number 81802101), the Development of Science and Technology, Jilin Province, China (Grant number2018010195JC), the Education Department of Jilin Province, China (Grant number JJKH20180239KJ), and the Health and Family Planning Commission of Jilin Province, China (Grant number 2017J074). The funders had no role in study design, data collection and analysis, decision to publish, or preparation of the manuscript.

## Acknowledgements

We thank the China Animal Health And Epidemiology Center (Qingdao, China) for their gifting of the serum samples and the whole bacterial antigen.

## References

1. Pappas G, Papadimitriou P, Akritidis N, Christou L, Tsianos EV. The new global map of human brucellosis. Lancet Infect Dis. 2006;6(2):91–9.

2. Goodwin ZI, Pascual DW. Brucellosis vaccines for livestock. Vet Immunol Immunopathol. 2016;181:51–58.

3. Mahajan V, Banga HS, Filia G, Gupta MP, Gupta K. Comparison of diagnostic tests for the detection of bovine brucellosis in the natural cases of abortion. Iran J Vet Res. 2017;18(3):183–9.

4. Paul S, Peddayelachagiri BV, Nagaraj S, Kingston JJ, Batra HV. Recombinant outer membrane protein 25c from Brucella abortus induces Th1 and Th2 mediated protection against Brucella abortus infection in mouse model. Mol Immunol. 2018;99:9–18.

5. Zheng WY, Wang Y, Zhang ZC, Yan F. Immunological characteristics of outer membrane protein omp31 of goat Brucella and its monoclonal antibody. Genet Mol Res. 2015;14(4):11965–74.

6. Soria-Guerra RE, Nieto-Gomez R, Govea-Alonso DO, Rosales-Mendoza S. An overview of bioinformatics tools for epitope prediction: implications on vaccine development. J Biomed Inform. 2015;53:405–14.

7. Greener JG, Sternberg MJ. Structure-based prediction of protein allostery. Curr Opin Struct Biol. 2018; 50:1–8.

8. Yin D, Li L, Song X, Li H, Wang J, Ju W, Qu X, Song D, Liu Y, Meng X, Cao H, Song W, Meng R, Liu J, Li J, Xu K. A novel multi-epitope recombined protein for diagnosis of human brucellosis. BMC Infect Dis. 2016;16:219.

9. Yin D, Li L, Song D, Liu Y, Ju W, Song X, Wang J, Pang B, Xu K, Li J. A novel recombinant multi-epitope protein against Brucella melitensis infection. Immunol Lett. 2016;175:1–7.

10. Simborio HL, Lee JJ, Bernardo Reyes AW, Hop HT, Arayan LT, Min W, Lee HJ, Yoo HS, Kim S. Evaluation of the combined use of the recombinant Brucella abortus Omp10, Omp19 and Omp28 proteins for the clinical diagnosis of bovine brucellosis. Microb Pathog. 2015; 83-84:41–6.

11. Xu J, Qiu Y, Cui M, Ke Y, Zhen Q, Yuan X, Yu Y, Du X, Yuan J, Song H, Wang Z, Gao G, Yu S, Wang Y, Huang L, Chen Z. Sustained and differential antibody responses to virulence proteins of Brucella melitensis during acute and chronic infections in human brucellosis. Eur J Clin Microbiol Infect Dis. 2013; 32(3):437–47.

12. Escalona E, Sáez D, Oñate A. Immunogenicity of a Multi-Epitope DNA Vaccine Encoding Epitopes from Cu-Zn Superoxide Dismutase and Open Reading Frames of Brucella abortus in Mice. Front Immunol. 2017; 8:125.

13. Clausse M, Díaz AG, Ibañez AE, Cassataro J, Giambartolomei GH, Estein SM. Evaluation of the efficacy of outer membrane protein 31 vaccine formulations for protection against Brucella canis in BALB/c mice. Clin Vaccine Immunol. 2014; 21(12):1689–94.

14. Risso GS, Carabajal MV, Bruno LA, Ibañez AE, Coria LM, Pasquevich KA, Lee SJ, McSorley SJ, Briones G, Cassataro J. U-Omp19 from Brucella abortus Is a Useful Adjuvant for Vaccine Formulations against Salmonella Infection in Mice. Front Immunol. 2017;8:171.

15. Solís García Del Pozo J, Lorente Ortuño S, Navarro E, Solera J. Detection of IgM antibrucella antibody in the absence of IgGs: a challenge for the clinical interpretation of brucella serology. PLoS Negl Trop Dis. 2014; 8(12):e3390.

16. Zygmunt MS, Diaz MA, Teixeira-Gomes AP, Cloeckaert A. Cloning, nucleotide sequence, and expression of the brucella melitensis sucb gene coding for an immunogenic dihydrolipoamide succinyltransferase homologous protein. Infect Immun. 2001;69:6537–40.

17. Hou Q, Sun X, Zhang J, Liu Y, Wang Y, Jin Z. Modeling the transmission dynamics of sheep brucellosis in Inner Mongolia Autonomous Region, China. Math Biosci. 2013;242(1):51–8.

18. Deqiu S, Donglou X, Jiming Y. Epidemiology and control of brucellosis in China. Vet Microbiol, 2002;90(1-4): 165–82.

19. Swets JA. Measuring the accuracy of diagnostic systems. Science. 1988;240:1285–93.

